# Tumor-Associated Macrophages Expand Chemoresistant, Ovarian Cancer Stem-Like Cells

**DOI:** 10.1101/2023.07.17.549067

**Authors:** Allison C Sharrow, Madeline Ho, Aakriti Dua, Raquel Buj, Kim R.M. Blenman, Sandra Orsulic, Ronald Buckanovich, Katherine M Aird, Lily Wu

## Abstract

The persistence of ovarian cancer stem-like cells (OvCSCs) after chemotherapy resistance has been implicated in relapse. However, the ability of these relatively quiescent cells to produce the robust tumor regrowth necessary for relapse remains an enigma. Since normal stem cells exist in a niche, and tumor-associated macrophages (TAMs) are the highest abundance immune cell within ovarian tumors, we hypothesized that TAMs may influence OvCSC proliferation. To test this, we optimized OvCSC enrichment by sphere culture and in vitro polarization of monocytes to a TAM-like M2 phenotype. Using cocultures that permitted the exchange of only soluble factors, we found that M2 macrophages increased the proliferation of sphere cells. Longer-term exposure (5-7 days) to soluble TAM factors led to retention of some stem cell features by OvCSCs but loss of others, suggesting that TAMs may support an intermediate stemness phenotype in OvCSCs. Although TAM coculture decreased the percentage of OvCSCs surviving chemotherapy, it increased the overall number. We therefore sought to determine the influence of this interaction on chemotherapy efficacy in vivo and found that inhibiting macrophages improved chemotherapy response. Comparing the gene expression changes in OvCSCs cocultured with TAMs to publicly available patient data identified 34 genes upregulated in OvCSCs by exposure to soluble TAM factors whose expression correlates with outcome. Overall, these data suggest that TAMs may influence OvCSC proliferation and impact therapeutic response.

## Introduction

Ovarian cancer is the fifth-deadliest cancer in US women, with approximately 65% of women with ovarian cancer dying as a result of this disease.^1^ Part of this high death rate is due to the fact that 70% of women are diagnosed at advanced stage (III or IV), when their cancer has already spread to distant sites.^2^ Another element in ovarian cancer’s high death rate is relapse. Although more than half of patients achieve complete response to first-line therapy (a combination of surgical debulking and paclitaxel and/or carboplatin chemotherapy), 75% of patients will relapse.^3^ Once patients relapse, their survival outcome worsens to a median survival of only 12-24 months.^3^ It is therefore crucial that we obtain a better understanding of the drivers of relapse to improve clinical outcomes for patients with ovarian cancer.

A large body of work has implicated a chemoresistant, stem-like subpopulation of cancer cells in ovarian cancer treatment failure.^4–9^ These ovarian cancer stem-like cells (OvCSCs) are more resistant to chemotherapy than the bulk tumor cells.^4, 10, 11^ This has been documented clinically by their relative enrichment in patient tumors after chemotherapy treatment.^11, 12^ OvCSCs are better able to grow tumors in murine models than bulk tumor cells, indicating that they possess the capacity to not only survive chemotherapy treatment, but also cause relapse.^6, 8^ Accordingly, worse clinical outcomes are observed in patients with high expression of OvCSC markers (including ALDH1, CD117, CD133, and CD44) in their untreated tumors and in patients with enrichment in these markers after chemotherapy treatment.^12, 13^ As such, better understanding the complex biology of OvCSCs has the potential to identify novel therapeutic approaches to prevent relapse.

One paradoxical aspect of OvCSCs is that, while they are felt to play a major role in driving disease recurrence, they are relatively quiescent compared to their non-stem counterparts.^4, 14^ Exactly how this relatively nonproliferative population of cells supports the robust tumor growth necessary to drive tumor recurrence remains unclear. One possibility is that surrounding host cells can increase the proliferative potential of OvCSC cells. Indeed, many normal adult stem cells exist in a niche composed of non-stem cells that support stem cell function.^15, 16^ We therefore hypothesized that components of the host microenvironment could stimulate OvCSC proliferation.

Tumor-associated macrophages (TAMs) are the predominant immune component of ovarian cancer in both solid tumors and ascites.^17, 18^ TAMs have been implicated in increased tumor angiogenesis and metastasis and may therefore impact patient survival.^19–21^ TAMs derive from normal monocytes that become reprogrammed by exposure to cancer cells. Normal macrophages are categorized along a spectrum as M0 (naïve), M1 (inflammatory), or M2 (immunosuppressive). Although these distinctions are not absolute, TAMs commonly express many of the M2 markers (CD163, CD206, low MHC class II molecules, interleukin 10, and [in mice] arginase 1) and are therefore thought to be largely protective against immune rejection of the tumor.^22^ TAMs have been reported to support the stemness of OvCSCs through secretion of specific cytokines that have known effects on the proliferation of bulk ovarian cancer cells.^23–28^ Although evidence suggests that TAMs influence OvCSCs, the direct effect of TAMs on OvCSC proliferation and treatment has only recently begun to be explored.

Given the abundance of TAMs in the tumor microenvironment, we sought to elucidate the influence of TAMs on OvCSC proliferation and treatment failure. After optimizing the in vitro enrichment of OvCSCs and the in vitro polarization of monocytes into a TAM-like M2 phenotype, we tested the impact of TAMs on OvCSC proliferation and found that coculture increased OvCSC proliferation in both the absence and presence of chemotherapy. Because of the documented effects of TAMs on ovarian cancer stemness when cultured over shorter timescales^23, 24^, we examined the effect of longer-term cocultures on the stem cell features of OvCSCs (adherence, chemoresistance, and OvCSC gene expression) and found augmented stem cell features in cocultured OvCSCs that may indicate an intermediate stem cell phenotype. We found a core set of 34 genes that were upregulated in OvCSCs cocultured with TAMs that correlate with worse progression-free and overall survival in patients. Finally, consistent with the TAM support of OvCSCs observed in vitro, targeting TAMs with CSF-1R inhibition improved response to chemotherapy in vivo. Combined, this work indicates a pro-proliferative role of TAMs on CSCs to increase chemoresistance, and further supports interactions between TAMs and OvCSCs as a clinical target in ovarian cancer.

## Materials and Method

### Cells and Adherent Cell Culture

ID8 and Kuramochi cells were obtained from Dr. Oliver Dorigo (Stanford University). ID8 cells were cultured in KnockOut Medium (Gibco) supplemented with 10% KnockOut Serum Replacement (Gibco), 2 mM L-glutamine (Gibco or Corning), 1X penicillin/streptomycin (Gibco or Corning), and 1% insulin-transferrin-selenium (Corning or Gibco). ID8 cultures were passaged using Accutase (StemCell Technologies or ThermoFisher), then counted and seeded. Kuramochi cells were cultured in RPMI (Corning or Gibco) supplemented with 10% heat-inactivated GenClone fetal bovine serum (HI-FBS, Genesee Scientific). Kuramochi cultures were passaged using Trypsin (Gibco or Corning), then counted and seeded.

### Sphere Culture

For sphere culture, ID8 and Kuramochi cells were dislodged from plastic as for passaging. They were then plated in ultra-low attachment flasks (Corning) at a density of 2×10^5^-3.3×10^5^ c/mL. Kuramochi and ID8 cells were plated in KnockOut Medium supplemented with 10% KnockOut Serum Replacement, 2 mM L-glutamine, 1X penicillin/streptomycin, 20 ng/mL fibroblast growth factor 2 (FGF2), and 20 ng/mL epidermal growth factor (EGF). ID8 cells were additionally supplemented with 1% insulin-transferrin-selenium. After 1 week, fresh medium was added in an amount equal to 50% of the starting volume. After 2 weeks of culture, spheres were used for experiments. For sphere digestion, Kuramochi spheres were incubated with trypsin, and ID8 spheres were incubated with Accumax (Stem Cell Technologies) at 37 °C with pipetting until spheres were dissociated. Cells were then counted and used for experiments.

### Sphere Chemoresistance

To compare the chemoresponsiveness of adherent and sphere cells, 2-week spheres and adherent cells were digested to obtain a single-cell suspension, counted, and plated at densities that would yield similar cell numbers at endpoint. Cell numbers plated into each well of a 96-well plate were as follows: Kuramochi spheres, 3.5×10^4^; Kuramochi adherent, 900; ID8 spheres, 5×10^3^; ID8 adherent, 750. Kuramochi cells were plated in triplicate with no drug and the following doses of paclitaxel (nM): 1, 2, 4, 5, 6, 8, 10, 25, 50, 100, and 1000 and the following doses of carboplatin (μM): 0.25, 0.5, 0.75, 1, 1.5, 2, 4, 6, 8, 10, 15, 20, 50, and 100 then incubated for 7 days. At endpoint, relative cell amounts were quantified using CellTiter Glo 3D (Promega) with the resulting luminescence measured with a CLARIOstar plate reader (BMG). ID8 were plated in triplicate with no drug and the following doses of paclitaxel (nM): 1, 2, 4, 6, 8, 10, 100, and 500 and the following doses of carboplatin: 1, 10, 20, 50, 75, 100, and 250 then incubated for 5 days. At endpoint, relative cell amounts were quantified using CellTiter 96 AQueous One Solution (Promega), and absorbance was measured with a CLARIOstar plate reader at 490 nm and 700 nm. Specific absorbance was calculated for each well as A_490 nm_ - A_700 nm_. Each dose was repeated in at least three independent experiments. Percent survival versus no drug control was calculated, outliers were removed based on Grubbs’ Test, normalcy was confirmed with the Shapiro-Wilk Test, and statistical significance was determined with ANOVA with post-hoc Tukey.

### In Vitro Macrophage Differentiation and Polarization

For human M1 and M2 polarization, CD14^+^ monocytes were isolated from healthy peripheral blood mononuclear cells (PBMCs) obtained from the UCLA CFAR Virology Core using human CD14 MicroBeads (Miltenyi Biotec) and plated at a density of 3.6×10^5^ cells/cm^2^ in RPMI supplemented with 2 mM L-glutamine, 1X penicillin/streptomycin, and 50 ng/mL recombinant human macrophage colony-stimulating factor (M-CSF, BioLegend). This was defined as Day 1. On Day 2, medium was changed to RPMI supplemented with 10% HI-FBS, 2 mM L-glutamine, 1X penicillin/streptomycin, and 50 ng/mL recombinant human M-CSF. On Day 4, medium was changed to “polarization medium” comprising RPMI supplemented with 10% HI-FBS, 2 mM L-glutamine, 1X penicillin/streptomycin, and appropriate human recombinant cytokines. M1 polarization cytokines included: 50 ng/mL M-CSF, 50 ng/mL granulocyte-macrophage colony-stimulating factor (GM-CSF, BioLegend), 50 ng/mL interferon γ (IFNγ, BioLegend), and 60 ng/mL lipopolysaccharide (LPS, Sigma). M2 polarization cytokines included: 50% Kuramochi sphere conditioned medium, 50 ng/mL M-CSF, and 30 ng/mL interleukin 4 (IL-4, Biolegend). On Day 7, cells were ready for use.

For murine M1 and M2 polarization, bone marrow was harvested from female C57BL/6 mice and plated at a density of 2.5×10^6^ cells/cm^2^ in DMEM (4.5 g/L glucose; Gibco or Corning) supplemented with 10% HI-FBS, 2 mM L-glutamine, 1X penicillin/streptomycin, and 50 ng/mL recombinant mouse M-CSF (BioLegend). This was defined as Day 1. On Day 4, medium was replaced with fresh medium of the same formulation. On Day 7, medium was changed to polarization medium containing DMEM (4.5 g/L glucose) supplemented with 10% HI-FBS, 2 mM L-glutamine, 1X penicillin/streptomycin, and appropriate murine recombinant cytokines. M1 polarization cytokines included: 50 ng/mL M-CSF, 50 ng/mL GM-CSF (BioLegend), 30 ng/mL IFNy (BioLegend), and 50 ng/mL LPS. M2 polarization cytokines included: 50% ID8 sphere conditioned medium, 50 ng/mL M-CSF, 50 ng/mL IL-4 (BioLegend), and 10 ng/mL interleukin-10 (IL-10, BioLegend). On Day 9, cells were ready for use.

To prepare conditioned medium, species-matched sphere cells (single-cell digestion) were plated at a density of 3.3×10^4^ cells/mL in ultra-low attachment flasks in the same medium that the macrophages would be polarized in (without cytokines). After 48 hours, medium was collected and filtered through a 0.22 μm filter. Culture medium was then aliquoted into single-use aliquots and frozen at −80 °C.

### Gene Expression Analysis

#### RT-PCR

Relative gene expression for OvCSC markers and macrophage polarization markers was measured using real-time RT-PCR. RNA was isolated using a silica-based column (RNeasy Kit [QIAGEN], Total RNA Miniprep Kit [BioPioneer], or Isolate II RNA Kit [Bioline]) and reverse transcribed with the High-Capacity Reverse Transcription Kit (Applied Biosystems). PCR reactions included PowerUP SYBR Green Master Mix (Applied Biosystems), 500 nM each primer (except human ALDH1A1, which included 300 nM each primer), and 1 ng cDNA (except murine ALDH1A3 and CD24, which used 10 ng/reaction and ALDH1A2 and CD133, which used 100 ng/reaction). Genes and primers are detailed in **Supplemental Table 1**. The thermocyclers used included Quantstudio 3 96-well and Quantstudio 5 384-well thermocyclers (Applied Biosystems). Data were analyzed in Applied Biosystem’s Relative Quantification App or calculated manually using the ΔΔCt method in Excel (Microsoft).

#### RNA-seq

Changes in global gene expression in cocultured spheres was measured using RNA-seq. For these cocultures, 6.9×10^6^ CD14^+^ cells from healthy human donors were plated on the top of 75-mm, 0.4-μm polycarbonate Transwell inserts (Costar #3419) and polarized to an M2 phenotype as described above. After polarization, 7.5×10^5^-10^6^ Kuramochi spheres were plated in each of two 100-mm Poly-HEMA–coated dishes (Poly-HEMA from Sigma) in fresh KnockOut medium without cytokines or growth factors. The Transwell chambers containing polarized M2 macrophages were added to one dish for each experiment. The other remained as monoculture. After 7 days, Transwell chambers were discarded, and sphere cells were collected into RNA later (Qiagen) and stored at −80 °C. Three independent experiments were performed, each with a different donor for the M2 macrophages. RNA was isolated from all samples in one batch using the Total RNA Miniprep Kit (BioPioneer). The UCLA Technology Center for Genomics & Bioinformatics performed library preparation and sequencing.

Libraries for RNA-Seq were prepared with the KAPA Stranded mRNA-Seq Kit. The workflow consisted of mRNA enrichment and fragmentation, first-strand cDNA synthesis using random priming followed by second-strand synthesis converting cDNA:RNA hybrids to double-stranded cDNA (dscDNA) with dUTP incorporation into the second cDNA strand. cDNA generation was followed by end repair to generate blunt ends, A-tailing, adaptor ligation, and PCR amplification. Different adaptors were used to multiplex samples in one lane. Sequencing was performed on an Illumina HiSeq 3000 for SE 1×50 run. Data quality check was performed on an Illumina SAV. Demultiplexing was performed with Illumina Bcl2fastq v2.19.1.403 software. Reads were mapped using STAR 2.27a, and read counts per gene were quantified using the human Ensembl GRCh38.97 GTF file.^29^ Read counts were normalized by CPM +1.0E-4 in Partek Flow (v7.0, Copyright 2019, Partek Inc., St. Louis, MO, USA). Differential gene expression was calculated using DESeq2. Pathway analysis was performed using Gene Set Enrichment Analysis (Broad Institute, Cambridge, MA, USA).

### Aldefluor Staining

To measure Aldefluor (Stem Cell Technologies^30^) fluorescence by flow cytometry, cells were removed from their culture vessels and digested into a single cell suspension as described in Cells and Adherent Cell Culture and Sphere Culture. Cells were pelleted and resuspended in Aldefluor buffer at a concentration of 10^6^ cells/mL. For each mL, 5 μL Aldefluor substrate was added. Cells were mixed by gentle vortexing, and 500 μL was immediately added to a second tube containing 5 μL DEAB. Cells were incubated for 30 min at 37 °C, then washed and stained with 3 μM DAPI and analyzed by flow cytometry on one of the following cytometers: FACS Aria (BD Biosciences), LSR II (BD Biosciences), or Fortessa (BD Biosciences). Data were analyzed using Diva (BD Biosciences) or FlowJo software (BD Biosciences). Cells were gated by forward and side scatter, doublets were excluded using forward and side scatter height by width plots, and viable cells were identified by DAPI exclusion. DEAB was used to set the Aldefluor^high^ gate with a background of 0.1%.

To measure Aldefluor fluorescence microscopically, cell culture supernatants were harvested, centrifuged, and stained as above. After washing, cells were deposited into cell culture chamber slides. Adherent cells were stained *in situ* through the addition of Aldefluor substrate diluted in buffer as for suspension staining. Dead cells were excluded through the addition if 0.3 μM DAPI prior to imaging. Aldefluor and DAPI fluorescence were captured with a Nikon Eclipse Ti2-E automated microscope with the same acquisition settings for all images. Because of the limited Aldefluor fluorescence generated by adherent cells, adherent cultures were permeabilized after Aldefluor imaging with the dropwise addition of ethanol while remaining on the automated microscope stage, then the same field was imaged again for DAPI to identify total cells in the well. This was not possible for spheres as the addition of ethanol would cause them to move within the well. All images were scaled the same for analysis. In FIJI, regions of interest (ROIs) encompassing spheres were drawn manually. Adherent ROIs were generated by creating masks of the DAPI-stained images and applying these ROIs to the Aldefluor image. Fluorescence histograms were obtained for all ROIs and aggregated. Frequencies were calculated from the count for each intensity divided by the total counts and multiplied by 100 to convert to a percentage.

For assessment of Aldefluor staining in cocultured cells, cells were additionally stained with 1.25 ng/mL Hoechst 33342 during the Aldefluor incubation to identify total cells without the need for subsequent permeabilization. Adherent cells were imaged using a MuviCyte (PerkinElmer). All images for a specific device were captured using the same acquisition settings, and all images were scaled the same before analysis. For the adherent fraction, total cell numbers were calculated in FIJI using the Hoechst images. Aldefluor thresholding was set using the matched DEAB controls with a background level of ∼0.5%. Using this thresholding, the number of Aldefluor^high^ cells was quantified in FIJI. To generate the percent Aldefluor^high^, the number of Aldefluor^high^ cells were divided by the total number of cells and multiplied by 100. Statistical significance was measured using a three-way ANOVA.

### Coculture

M2 macrophages were polarized on the upper surface of 0.4-μm cell culture inserts (Corning and CELLTREAT). Digested spheres were plated into the tissue culture-treated wells below. Experiments used 24-well plates with 2.5×10^4^ Kuramochi sphere cells per well and 2.0×10^4^ ID8 sphere cells per well. When chemotherapy drugs were included, Kuramochi spheres were treated with 75 nM paclitaxel and 75 μM carboplatin, and ID8 spheres were treated with 250 nM paclitaxel and 50 μM carboplatin.

Proliferation endpoints were measured using CellTiter 96 AQueous One Solution after 7 days for Kuramochi and 5 days for ID8. Absorbance was measured with a CLARIOstar plate reader at 490 nm and 700 nm. Specific absorbance was calculated for each well as A_490 nm_ - A_700 nm_. Outliers were removed based on Grubbs’ Test. Normalcy was confirmed with the Shapiro-Wilk Test. Statistical significance was calculated using paired T tests and corrected with the Holm-Bonferroni method.

For growth curves, one coculture set was plated for each timepoint to be analyzed. Cell amounts were quantified at 24-hour intervals over 7 days for Kuramochi cells and 5 days for ID8 cells using CellTiter 96 AQueous One Solution and read and analyzed as above.

For adherence, cocultures were analyzed at 24 hours and 7 days. Cells were fixed with 4% paraformaldehyde (Electron Microscopy Sciences) for 15 min then stained with 0.5% crystal violet (ThermoFisher) in 20% methanol (ThermoFisher) for 5 min. Cells were then washed thrice with water and air-dried. Whole wells were imaged using a Nikon Eclipse Ti2-E automated microscope. The surface area covered by crystal violet-stained cells was quantified in FIJI. Fold-change was calculated as Area-_coculture_/Area_monoculture_. Outliers were removed based on Grubbs’ Test. Normalcy was confirmed with the Shapiro-Wilk Test. Statistical significance was calculated using paired T Tests.

### In Vivo Treatment

To stably mark ID8 for bioluminescence imaging, the firefly luciferase gene was cloned out of the pGL3-basic plasmid (Promega) using primers that added a BamHI site at the 5’ end and an XhoI site at the 3’ end, removing the stop codon. This product was then cloned in-frame 5’ of mStrawberry, producing a fusion product driven by a CMV promoter (**Supplemental Figure 1**). The resulting plasmid was transfected into 293T cells along with second-generation lentivirus packaging plasmids VSV-G and 8.2 using calcium phosphate transfection. After two days, virus-containing culture supernatant was harvested and applied to adherent ID8 cells at high titer to ensure transfection of the slowly proliferating Aldefluor^high^ cells. This was confirmed by flow cytometry using the Aldefluor staining procedure described above. Successfully transfected, mStrawberry^+^ cells were sorted on a FACS Aria, and firefly luciferase activity was confirmed in vitro prior to implantation.

From these adherent cells, spheres were generated as described above. Single-cell suspensions of adherent and sphere ID8 cells were obtained and mixed in a ratio of 90% adherent cells and 10% sphere cells. Cells were mixed with Matrigel (Corning) at a final concentration of 5 mg/mL. A total of 10^6^ cells were injected intraperitoneally (IP) into each 6-week-old C57BL/6 female mouse. Beginning at week 6 after cell injection, mice underwent weekly bioluminescence imaging to monitor tumor growth. At week 10, tumors were readily detectable in all mice, and mice were stratified into four treatment groups such that each group had similar means, medians, total fluxes, and standard deviations. In addition, a series of Two-Sample Kolmogorov-Smirnov Tests were performed to confirm that all groups derived from the same distributions. A random number generator was then used to assign treatments to each group of animals.

Each treatment group contained 7 mice and received one of the following treatments: (1) vehicle control; (2) paclitaxel only, 20 mg/kg IP weekly; (3) GW2580 only, 160 mg/kg 5 days per week by oral gavage; and (4) paclitaxel + GW2580, same dosing as for monotherapies. Mice were treated for 4.5 weeks. Paclitaxel (Sigma) was reconstituted in 50% cremophor (Sigma) in ethanol (ThermoFisher) then diluted 50-fold into 0.9% sodium chloride injection solution (Hospira) just prior to injection. The macrophage colony-stimulating factor 1 receptor (CSF-1R) inhibitor GW2580 was reconstituted in 0.1% hydroxypropyl methyl cellulose (Sigma) and 0.5% Tween 20 (ThermoFisher) in water. For each treatment, mice were weighed, and the appropriate doses of each drug were administered based on weight.

For bioluminescence imaging, mice were anesthetized with 2.5% isoflurane, shaved, and injected retro-orbitally with 100 μL of 30 mg/mL luciferin. Mice were imaged weekly with an IVIS in vivo imaging system (PerkinElmer) using consistent settings throughout the experiment. Throughout the treatment period, ascites was removed via paracentesis as needed. Mice were anesthetized with 2.5% isoflurane and ascites fluid was carefully removed using a 3-mL syringe with a 27G, ½-inch needle inserted into the abdominal cavity. The removed volume was carefully measured. At endpoint, mice underwent a final bioluminescence imaging and were then euthanized via cervical dislocation under anesthesia. Skin was blunt dissected away from the underlying muscle wall, and any ascites fluid was withdrawn and measured. The abdominal and thoracic cavities were opened with the mice pinned to dissecting trays. Bioluminescence images were used to identify and excise all tumor tissues. Total tumor weight was measured.

For analyses, outliers were excluded based on Grubbs’ Test, and normalcy was assessed using the Shapiro-Wilk Test. Differences in ascites volume were determined with the Kruskal-Wallis Test. Tumor weights were assessed with an ANOVA with post hoc Tukey. The proportion of mice showing reduced bioluminescence at endpoint vs treatment initiation was assessed with a Fisher’s Exact Test.

## Results

### Sphere Culture Enriches for Chemoresistant Ovarian Cancer Stem-Like Cells

To enrich for chemoresistant OvCSCs, Kuramochi and ID8 cells were cultured under nonadherent sphere-formation conditions for 2 weeks in ultra-low attachment flasks in KnockOut Medium supplemented with 20 ng/mL FGF2 and EGF. At the end of coculture, few single cells remained, with the majority being in dense, multicellular spheres that could become quite large and often aggregated (**Figure 1A**). At endpoint, the cell numbers decreased by 81% in Kuramochi and 42% in ID8 (**Figure 1B**), suggesting a loss of cells whose survival is not supported by nonadherent culture, such as bulk ovarian cancer cells. Consistently, we observed an increase in OvCSC markers *ALDH1A1*, *ALDH1A3*, and *CD24* in Kuramochi, and *ALDH1A1*, *CD44*, and C*D133* in ID8 (**Figure 1C, Supplementary Figure 2**). Supporting a role for OvCSCs in chemotherapy resistance, expression of these markers was similar to attached cells treated with the standard-of-care therapies paclitaxel and carboplatin (**Figure 1C**), which are known to enrich for OvCSCs.^31, 32^ Consistent with their higher expression of ALDH1 family members, spheres displayed higher Aldefluor fluorescence than their adherent counterparts (**Figure 1D**). A functional hallmark of OvCSCs is their chemoresistance. Indeed, spheres were more resistant to ovarian cancer standard-of-care therapies paclitaxel and carboplatin (**Figure 1E**). Together, these data suggest that sphere culture enriches for chemoresistant OvCSCs consistent with published literature supporting this enrichment method.^33, 34^

**Figure 1:**
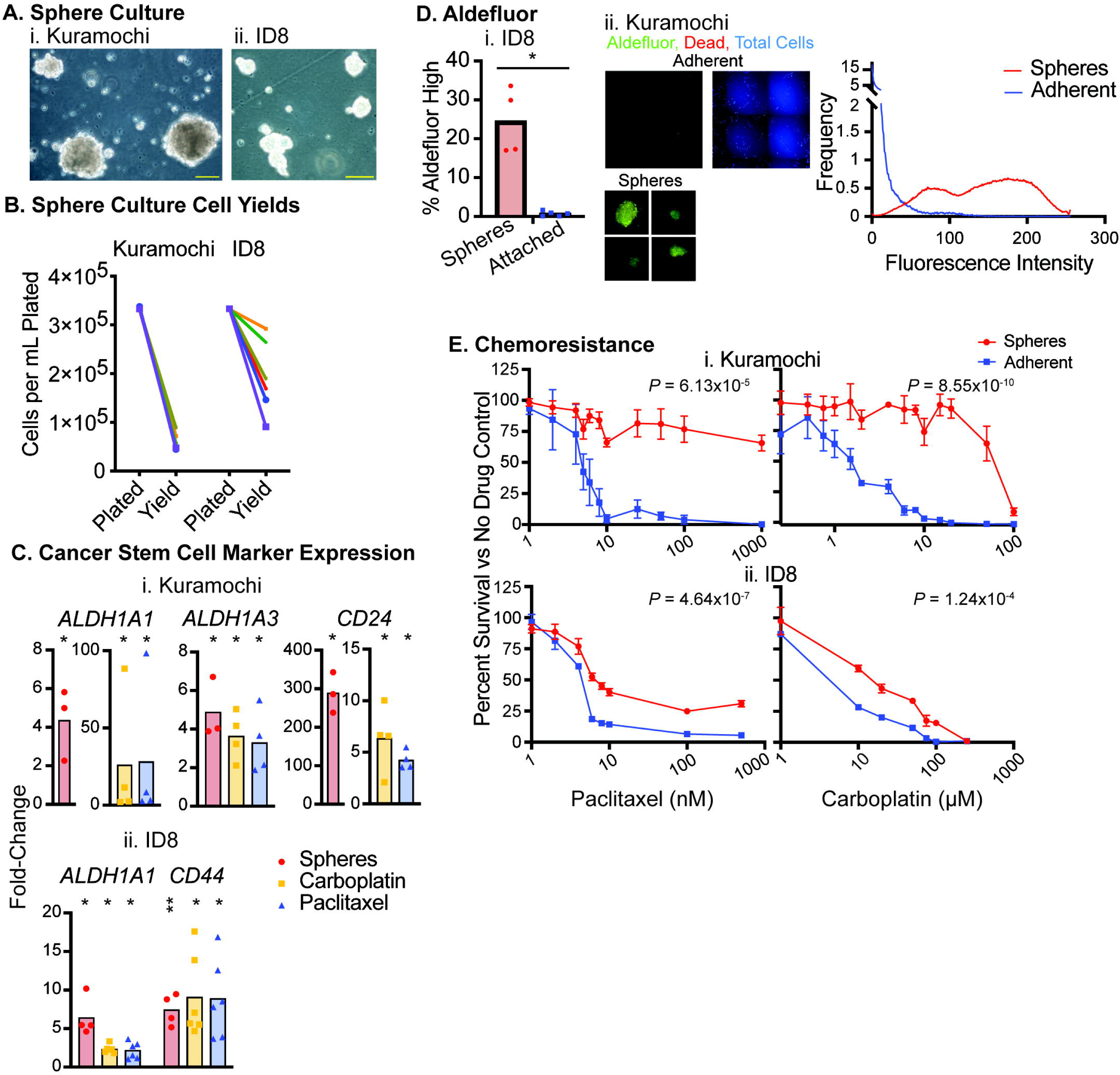
Sphere Culture Enriches for Chemoresistant Ovarian Cancer Stem-Like Cells. **(A)** Representative microscopic images of **(i)** Kuramochi and **(ii)** ID8 cells cultured for 2 weeks under nonadherent conditions with 20 ng/mL EGF and FGF2. Scale bars are 20 μm long. **(B)** Plots show the numbers of Kuramochi and ID8 cells per mL plated at the initiation of sphere culture (“Plated”) and the cell numbers per mL resulting from 2 weeks of sphere culture (“Yield”). Individual cultures (*n* = 6 per cell line) are joined by different colored lines. **(C)** Real-time RT-PCR measurement of expression of the indicated genes in **(i)** Kuramochi and **(ii)** ID8 cells. Y axis shows fold-change in: spheres vs attached (red), carboplatin-treated vs untreated attached (yellow) and paclitaxel-treated vs untreated attached (blue). Shaded bars show mean fold-change, and individual points show the results of independent biological replicates. Based on results of Shapiro-Wilk testing for normality, unmatched samples (spheres vs attached) were compared with their control using a t-test. Matched samples (carboplatin-treated / paclitaxel-treated vs untreated) were compared to their controls using a Mann Whitney Test. Statistical significance for each is presented with the following indicators: 0.001 ‘**’ 0.01 ‘*’ 0.05. **(D)** Aldefluor staining of spheres and attached cells. **(i)** ID8 cells were detected and quantified with flow cytometry and are presented as the percentage of Aldefluor high cells set using DEAB as a negative control. Shaded bars show mean, and individual points show the results of independent biological replicates. Statistical testing used a T test and is presented with the following indicator: 0.01 ‘*’ 0.05. **(ii)** Kuramochi cells were detected with microscopy and quantified using FIJI. Representative images are shown with Aldefluor in green and DAPI (dead cells) false colored red. For the adherent field, cells were also permeabilized and stained with DAPI to show total cells, which is colored in blue. Fluorescence intensity histograms of cells were generated in FIJI. Data are presented as frequency (percent of total counts) vs fluorescence intensity. **(E)** Chemoresistance of **(i)** Kuramochi and **(ii)** ID8 cells grown as spheres or adherent cultures in the indicated doses of paclitaxel (left) or carboplatin (right). Cell numbers were measured using CellTiter Glo for Kuramochi or CellTiter 96 AQueous One Solution for ID8 and normalized to matched untreated control. Mean and SEM of aggregated biological replicates are plotted. Statistical significance was determined with ANOVA with post-hoc Tukey and presented with the following indicators: 0 ‘***’ 0.001 ‘**’ 0.01 ‘*’ 0.05.

### Coculture of Spheres with M2 Macrophages Increases Their Proliferation

Cells in sphere culture proliferate only very slowly. To determine if TAMs could influence the proliferation of these cells, we generated TAM-like M2 macrophages in vitro. Primary monocytes were isolated from human PBMCs or murine bone marrow then polarized to an M2 phenotype, exhibiting gene expression consistent with TAMs found in human ovarian tumors (**Supplementary Figure 3**).^22^ In vitro-differentiated M2 macrophages were then cocultured with dissociated sphere cells separated by 0.4-μm cell culture inserts. As such, soluble mediators could pass through the membrane, but cells could not. Coculture of sphere cells with M2 macrophages increased sphere cell proliferation (**Figure 2A**). We then performed RNA-seq of cocultured and monocultured spheres. Consistent with the in vitro proliferative changes, Gene Set Enrichment Analysis of RNA-Seq data showed enrichment in multiple proliferation pathways in Kuramochi spheres cocultured with M2 macrophages compared to monocultured spheres, including NFκB, mTor, and MAP kinase (**Figure 2B**, **Table 1**). These data indicate that M2 macrophages increase the proliferation of OvCSCs.

**Figure 2:**
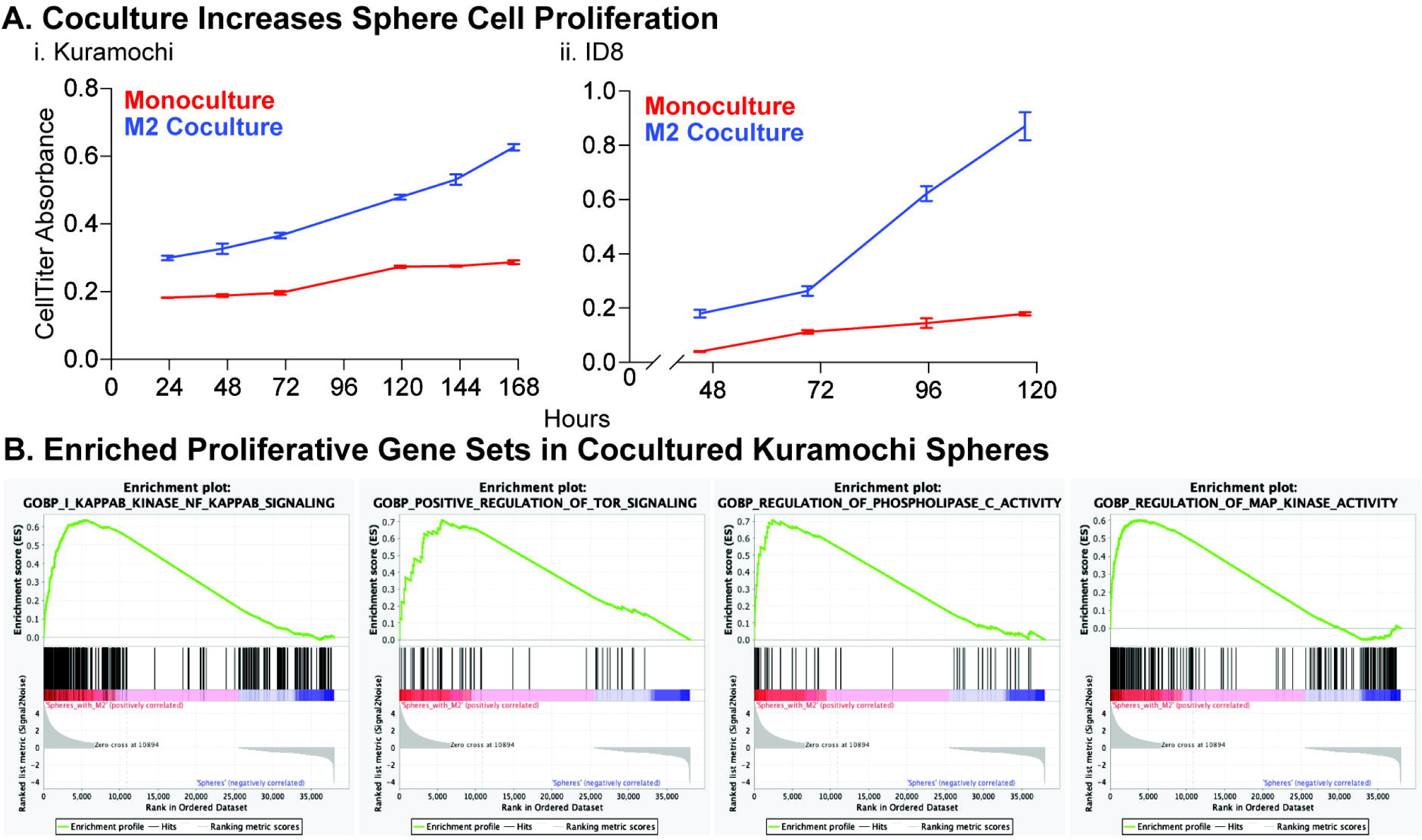
M2 Macrophages Increase OvCSC Proliferation. **(A) (i)** Kuramochi and **(ii)** ID8 sphere cells were cultured in monoculture or cocultured with M2 macrophages separated by 0.4 μm cell culture inserts. At the stated times after plating, independent cultures were analyzed with CellTiter 96 AQueous One Solution. Absorbance was calculated as the specific absorbance at 490 nm minus the background absorbance/scatter at 700 nm and plated vs time. Error bars show the SEM of technical replicates. **(B)** Enrichment plots from GSEA analysis of Kuramochi spheres cocultured with M2 macrophages for 7 days vs monocultured spheres.

### M2 Macrophages Supports an Intermediate Stemness Phenotype

Published literature has explored the impact of TAMs on OvCSC stemness over relatively short timescales.^23, 24^ We therefore sought to examine the the stem cell phenotype in OvCSCs after longer-term (5-7 days) coculture. As shown in **Figure 1** and published literature, nonadherent sphere cells display a chemoresistant, OvCSC phenotype that includes expression of stem cell markers.^34^ In contrast, adherent cells are relatively chemosensitive and have lower expression of ovarian cancer stem cell markers, consistent with a more differentiated phenotype.^34^ As such, cellular adhesion may serve as a phenotypic indicator of stemness. Coculture with M2 macrophages increased the number of sphere cells adhering to the tissue culture surface (**Figure 3A**). One possible explanation for this observation is that an overall greater number of cells are already present at this 24-hour timepoint and that the same proportion of cells is adhering. To test this, the adherence of Kuramochi spheres was compared between 7 days of monoculture and 24 hours of coculture — conditions that produce comparable numbers of cells (**Figure 3Bi**). In conditions with similar numbers of cells, the increased attachment due to coculture persisted, suggesting that TAMs alter the adherence of OvCSCs (**Figure 3Bii**). GSEA analysis identified 9 datasets enriched in cocultured Kuramochi spheres at an FDR < 0.05 related to cellular adhesion; all unique datasets with FDR q-values below 0.01 are presented in **Figure 3C**.

**Figure 3:**
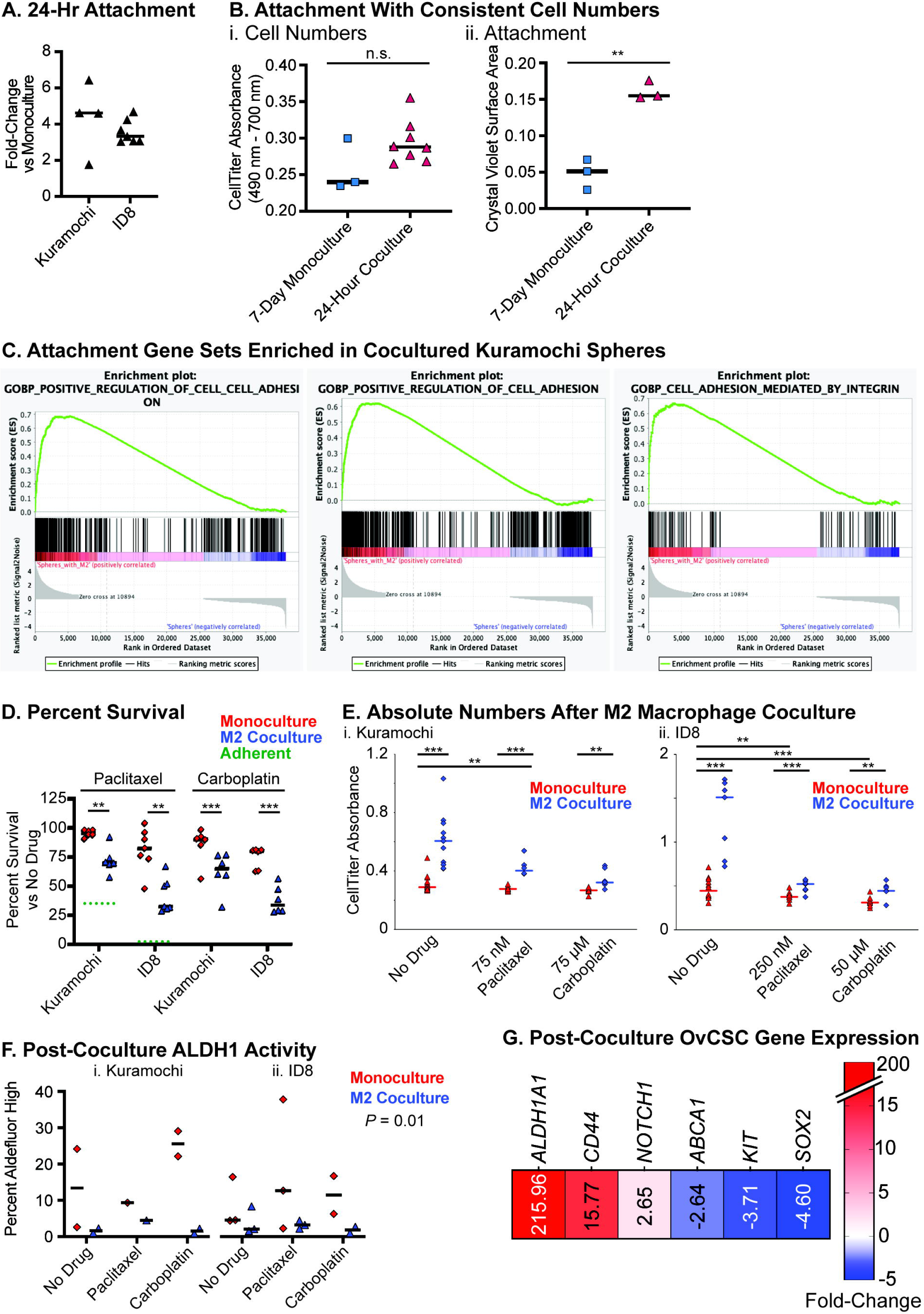
M2 Macrophages Support An Intermediate Stemness Phenotype. **(A)** Kuramochi and ID8 spheres were cultured for 24 hours in monoculture or coculture with M2 macrophages separated by 0.4 μm permeable cell culture inserts. After 24 hours, cells were fixed and stained with crystal violet. Whole wells were imaged and the area covered by cells was calculated using FIJI. Fold-change was calculated as cocultured vs monocultured spheres. Individual experiment datapoints are plotted with lines indicating medians. Statistical comparison between monoculture and M2 coculture was tested using paired T tests and found to be significant (Kuramochi: *P* = 0.02, ID8: *P* = 0.03). **(B)** To determine if the change in attachment was due to differences in cell numbers, 24-hour coculture was compared to 7-day monoculture in Kuramochi spheres, cultured as in (A). **(i)** CellTiter 96 AQueous One Solution was used to determine the relative cell numbers at these timepoints. Absorbance was calculated as the specific absorbance at 490 nm minus the background absorbance/scatter at 700 nm. Individual experiment datapoints are plotted with lines indicating medians. T tests were used to assess statistical difference. **(ii)** At the indicated timepoints, cells were fixed and stained with crystal violet. Whole wells were imaged, and the area covered by cells was calculated using FIJI. Individual experiment datapoints are plotted with lines indicating medians. T tests were used to assess statistical difference. **: *P* < 0.01. **(C)** Enrichment plots from GSEA analysis of Kuramochi spheres cocultured with M2 macrophages for 7 days vs monocultured spheres. **(D) (i)** Kuramochi and **(ii)** ID8 sphere cells were cocultured with M2 macrophages separated by 0.4 μm permeable cell culture inserts. After 7 days for Kuramochi and 5 days for ID8, the resulting sphere cells were measured using CellTiter 96 AQueous One Solution. Absorbance was calculated as the specific absorbance at 490 nm minus the background absorbance/scatter at 700 nm, then percent survival vs matched no drug control was calculated. Results of independent biological replicates are plotted with lines indicating medians. Red triangles indicate sphere monocultures, blue diamonds indicate M2 cocultures. Green dotted lines indicate survival of monoculture attached cells at 75 μM paclitaxel for Kuramochi and 30 nM paclitaxel for ID8 (ID8 spheres were cultured in 250 nM paclitaxel). Differences in survival between matched monoculture and M2 coculture samples were calculated using paired T tests, and significance threshold was adjusted for multiple comparisons using the Holm-Bonferroni method with α < 0.05. Black bars indicate significant differences and are indicated as: 0 ‘***’ 0.001 ‘**’ 0.01. **(E)** Absolute cell numbers of experiments from D are presented for **(i)** Kuramochi and **(ii)** ID8 sphere cells. Individual biological replicates are plotted with lines indicating medians. Statistical significance was calculated using paired T tests and corrected with the Holm-Bonferroni method. For each drug condition, monoculture was compared to coculture with M2 macrophages, and all conditions were compared to no drug monoculture. Black bars indicate statistically significant differences with α < 0.05. *P* values and cutoffs are detailed in **Supplemental Table 2** and indicated as: 0 ‘***’ 0.001 ‘**’ 0.01 ‘*’ 0.05. Red triangles indicate sphere monocultures, blue diamonds indicate M2 cocultures. **(F)** At coculture endpoint, cells were stained with Aldefluor and Hoeschst then imaged. Percent Aldefluor-high cells was calculated in FIJI using matched DEAB controls to set thresholds. Differences were calculated with a 3-way ANOVA, finding a significant effect of macrophages (*P* = 0.01). **(G)** Kuramochi spheres were cultured in nonadherent conditions either in monoculture or cocultured with M2 macrophages separated by 0.4 μm cell culture inserts. After 7 days, spheres were isolated and underwent RNA-seq. Differentially regulated (FDR < 0.05) OvCSC genes in cocultured vs monocultured Kuramochi spheres are presented in a heatmap with their fold-change.

Because attachment is a feature of sphere cell differentiation, and sphere cells are more chemoresistant that adherent cells, the percent survival of monocultured and M2 cocultured spheres were compared. Coculture with M2 macrophages decreased the percentage of sphere cells surviving paclitaxel and carboplatin treatment (**Figure 3D**). However, this reduction in percent survival did not reach the survival level of adherent cells (green dotted lines in **Figure 3D**), nor did coculture consistently alter the percent survival in adherent cells compared to monoculture. Furthermore, absolute numbers showed that coculture increased the number of sphere cells surviving chemotherapy compared to monoculture, indicating that TAMs could increase OvCSC persistence after chemotherapy despite reducing the percent survival (**Figure 3E**). We further assessed stemness by measuring ALDH1 activity using Aldefluor staining and microscopy. Coculture reduced the percentage of Aldefluor^high^ cells compared to monoculture (**Figure 3F**). Analysis of RNA-seq data comparing cocultured to monocultured Kuramochi spheres found a mixture of upregulated and downregulated ovarian cancer stem cell genes (**Figure 3G**), and gene sets related to stem cell differentiation and stem cell developmental pathways (e.g., Notch, Wnt/β-catenin, FGF2, and EGF) were not enriched in cocultured cells. Taken together, these data suggest that long-term exposure to soluble factors from M2 macrophages may support an intermediate level of stemness in OvCSCs with regards to chemoresistance, percentage of Aldefluor^high^ cells, and expression of OvCSC genes.

### Macrophage Inhibition Improves Response to Paclitaxel

Because TAMs increase the numbers of OvCSCs surviving chemotherapy treatment (**Figure 3E**), we examined the effect of TAMs on chemotherapy response in vivo. To do so, ID8 cells were stably transduced to express firefly luciferase fused to mStrawberry at high titers to ensure transduction of the less proliferative OvCSCs. No differences were found in Aldefluor staining between the 85% of cells that were mStrawberry and firefly luciferase positive and the remainder that were negative (**Figure 4A**), suggesting that OvCSCs were transduced at the same rate as bulk cells. After sorting for mStrawberry^+^ cells, a mixture of 90% adherent and 10% sphere cells was injected IP into C57BL/6 mice. After tumor establishment, mice were treated with paclitaxel and the CSF-1R inhibitor GW2580 (which depletes macrophages) alone and in combination. Similar to previous reports, GW2580 reduced ascites volume independent of paclitaxel treatment (**Figure 4B**).^35^ At endpoint, bioluminescence imaging was used to identify and excise tumor nodules throughout the abdominal cavity, which were then weighed.

**Figure 4:**
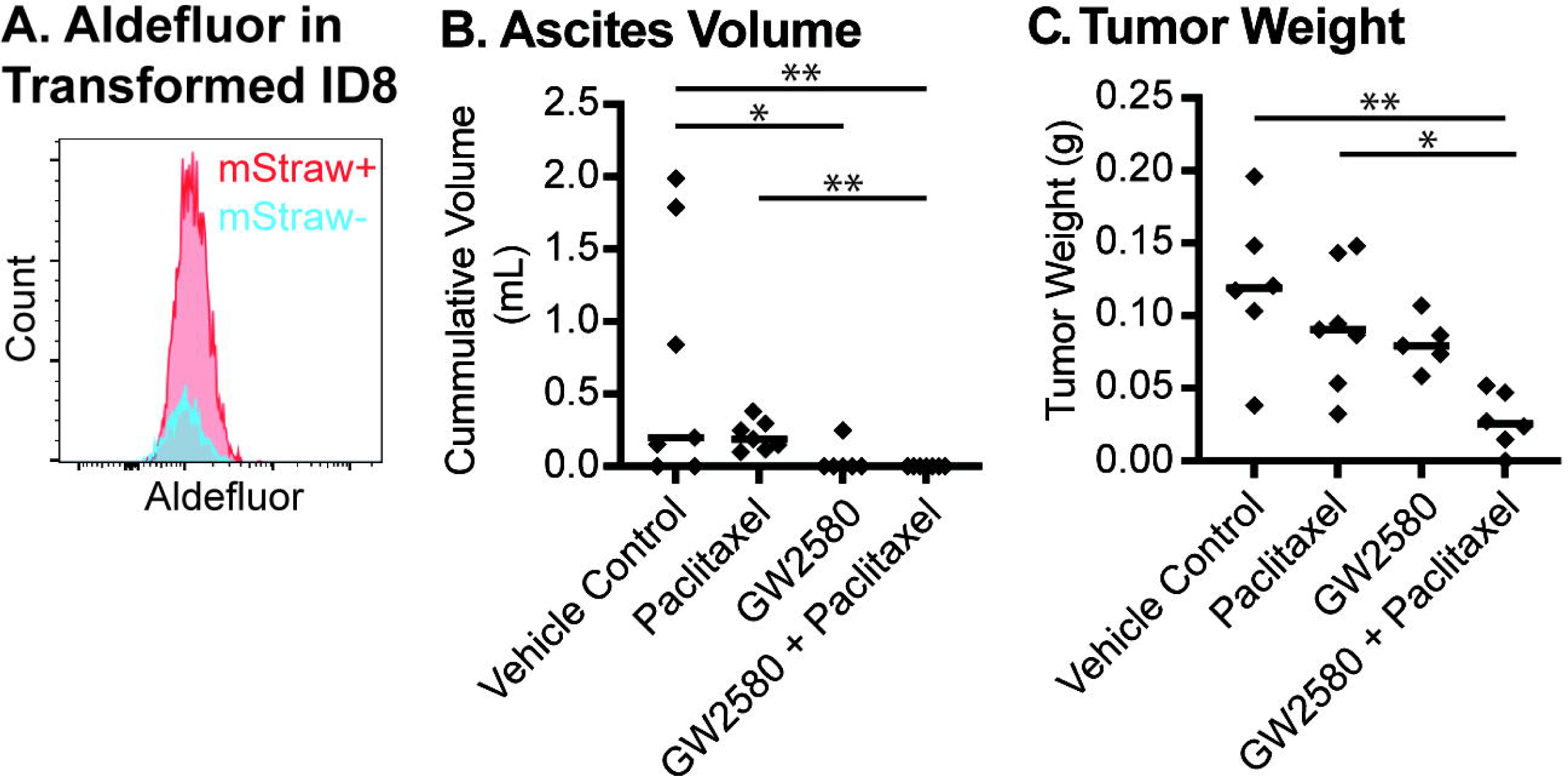
Macrophage Blockade Improves Chemotherapy Response. **(A)** ID8 cells were transduced with lentivirus containing firefly luciferase and mStrawberry. Cells were stained with Aldefluor, which was compared in transduced (mStraw^+^) and nontransduced (mStraw^-^) cells. **(B)** Throughout the treatment period and at endpoint, ascites was removed from the abdominal cavity with a syringe and needle and measured. Cumulative values are plotted for each treatment group: vehicle control, paclitaxel (20 mg/kg IP weekly for 5 weeks), GW2580 (160 mg/kg 5 days per week by oral gavage for 4 weeks 2 days), paclitaxel + GW2580 (same dosing as for monotherapies). Differences between groups were measured using a Kruskal-Wallis rank sum test for multiple independent samples with post-hoc Holm FWER-adjusted Conover *P*-values for all possible pairwise comparisons. Statistically significant differences (α < 0.05) are indicated with lines and the following identifiers: 0.001 ‘**’ 0.01 ‘*’ 0.05. **(C)** At endpoint, tumors were excised and weighed. Tumor weight is plotted for each treatment group (same as in B). Differences between groups were measured with an ANOVA with post-hoc Tukey comparing all possible pairwise comparisons. Statistically significant differences are indicated with lines and the following identifiers: 0.001 ‘**’ 0.01 ‘*’ 0.05.

Macrophage blockade successfully improved response to chemotherapy, yielding a significantly decreased tumor weight compared to untreated or paclitaxel-treated tumors (**Figure 4C**). Furthermore, combination therapy was the only treatment that reduced tumor burden below that observed during pretreatment imaging (4 of 7 vs 0 of 7 in all other groups, *P* = 0.015). Overall, macrophage blockade improved chemotherapy response by reducing ascites volume, tumor weight, and relative tumor burden.

To assess the clinically relevant genes altered during this interaction, RNA-seq data from cocultured Kuramochi spheres were compared to TCGA data. TCGA RNA-seq data were stratified by patients with poor outcome (less than 2 years survival) and good outcome (greater than 5 years survival). The resulting differentially expressed genes were compared to those of cocultured Kuramochi spheres vs monoculture spheres, and 50 of the genes upregulated in the poor outcome group were also upregulated in cocultured Kuramochi spheres (**Supplemental Table 3**). This 50-gene list generated from TCGA served as a training dataset that was then validated by testing for impacts on progression-free and overall survival using Gene Expression Omnibus datasets in KMplot.^36^ Greater expression of these 50 genes was found to correlate with worse outcomes (**Figure 5A**). Each of the genes in the 50-gene training list was then independently assessed for clinical correlation to progression-free survival, with 34 independently predicting outcome. (**Supplemental Table 3**). Only 9 of the 34 genes have been previously linked to outcome (see **Suppmental Table 3 for references**). Taken together, this refined, validated list has a more robust impact on progression-free survival (**Figure 5B**), with patients with high mean expression of these 34 genes surviving an average of 13 months shorter than patients with low mean expression of these genes (low-expression median survival: 50.03 months, high-expression median survival: 36.83 months). This suggests that a core group of genes that is altered in OvCSCs by exposure to TAMs correlates with worse clinical outcome.

**Figure 5:**
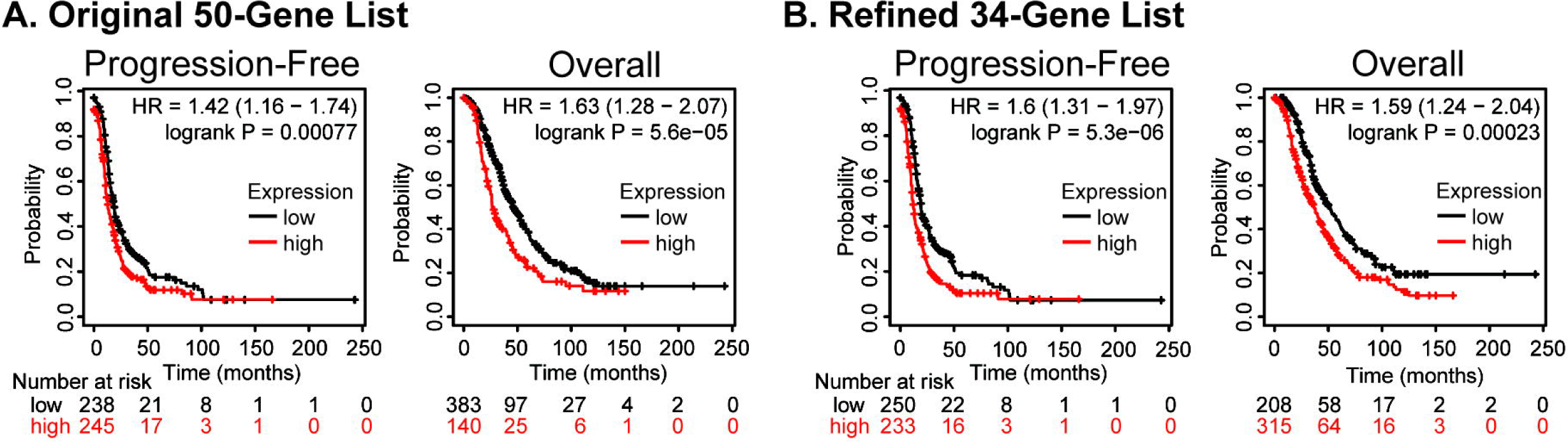
Common Gene Set Correlates with Worse Progression-Free and Overall Survival. Kaplan-Meyer curves generated using KMplot for the mean expression of the **(A)** initial list of 50 genes identified in the training set (TCGA) and the **(B)** 35 genes that remained significant in the validation set (KMplot GEO data sets) with HR > 1.2 and logrank *P* value < 0.05.

## Discussion

The subpopulation of cancer cells frequently termed OvCSCs have repeatedly been found to be relatively chemoresistant and able to regrow tumors, implicating them in ovarian cancer relapse.^4, 6, 8, 10, 11^ How these relatively quiescent cells are capable of rapidly regenerating tumors is an important question with serious clinical implications.^4, 14, 37^ Prior work implicated TAMs in stimulating the proliferation of bulk ovarian cancer cells.^38, 39^ We found that TAMs similarly increase the proliferation of OvCSCs and augment their stem cell phenotype. Importantly, we found that disrupting TAMs in vivo improved chemotherapy response.

Our coculture experiments established to model the interactions between OvCSCs and TAMs were specifically designed to avoid direct contact because the effects of direct contact have already been skillfully explored^23^, and we reasoned that soluble factors may potentially impact a greater number of OvCSCs. Despite coculture increasing the numbers of cells remaining after drug treatment, a lower percentage of cells survived. It was also observed that coculture increased the adherence of OvCSCs. Both of these phenotypes are associated with reduced stemness.^4, 6, 9, 33^ In keeping with this, coculture reduced a subset of OvCSC markers, including ALDH1 activity. Prior published works have shown TAM support for a stem cell phenotype, increasing stemness with short-term exposure to soluble TAM factors in several cancer types.^24, 40, 41^ In ovarian cancer, this is mediated by IL-8 signaling, which was similarly enriched in the GSEA analysis of cocultured Kuramochi spheres.^24^ Direct contact under intermediate timescales similarly favors an OvCSC phenotype, but through IL-6 signaling and *WNT5B*.^23^ GSEA analysis of cocultured Kuramochi spheres showed enrichment in IL-6 signaling and upregulation of *WNT5B*. Together, these data suggest that short-term or close contact between OvCSCs and TAMs may increase the OvCSCs within a tumor; but as the OvCSCs proliferate and move away from the TAMs, the effect of soluble factors may become more prominent. Our data suggest that sustained exposure to these soluble TAM factors supports an intermediate phenotype. This would mimic the processes that occur in healthy tissues in which a stem cell proliferates and forms transit amplifying cells that in turn differentiate. The presence of macrophages may therefore have a dual function of maintaining a stem cell pool while also favoring the expansion of a transit-amplifying–like cell. Future work is needed to more fully characterize the presence of transit-amplifying–like cells in ovarian cancer.

This work adds to the growing body of literature implicating TAMs in tumor progression and chemoresistance. Our findings, consistent with that of others suggests that targeting TAMs can increase chemotherapy response in ovarian cancer. While many studies targeting TAMs focus on the immune compartment, this work would support investigating the impact of TAM-targeting drugs on OvCSCs.

## Acknowledgements

This work was supported by the following grants: CDMRP Ovarian Cancer Research Program W81XWH-17-1-0160 (Wu), UCLA Tumor Immunology Training Grant 4T32CA009120-40 (Sharrow), UCLA Interdisciplinary Training in Virology and Gene Therapy 5T32AI060567-14 (Sharrow), Tobacco-Related Disease Research Program Postdoctoral Fellowship 27FT-0023 (Sharrow), Iris Cantor Women’s Health Center / CTSI Young Investigator Award (Sharrow), NIH 5P30 AI028697 (UCLA CFAR), and NIH P30 CA016042 (UCLA Flow Cytometry Core).

## Supplementary Figures

**Supplemental Figure 1: Plasmid Map of Firefly Luciferase mStrawberry Construct**

Map was created using SnapGene’s Restriction and Insertion Cloning tool.

**Supplemental Figure 2: Expression of Ovarian Cancer Stem-Like Cell Markers in Spheres and Treated Adherent Cells**

Real-time RT-PCR measurement of expression of the indicated genes. Y axis shows fold-change in **(i)** Kuramochi and **(ii)** ID8 spheres vs attached (red), carboplatin-treated vs untreated attached (yellow), and paclitaxel-treated vs untreated attached (blue).

Shaded bars show means, and individual data points are the results from independent biological replicates. Unmatched samples (spheres vs attached) were compared with their control using a t-test. Matched samples (carboplatin-treated / paclitaxel-treated vs untreated) were compared to their controls using a Mann Whitney Test. Statistical significance for each is presented with the following indicators: 0.001 ‘**’ 0.01 ‘*’ 0.05.

**Supplemental Figure 3: In Vitro Polarization of Macrophages**

Expression of the listed genes was measured for in vitro-differentiated M1 and M2 macrophages using real-time RT-PCR. Expression was calculated using the ΔΔCt method. Gene expression was also measured in isolated human TAMs and expressed as fold-change for TAMs vs in vitro-differentiated M1 macrophages. # indicate that the gene was not tested for that species. M2 macrophages and TAMs failed to express IL-12; therefore, these data are not plotted. Differences were assessed with Mann-Whitney tests, with significance indicated as follows: ‘***’ 0.001 ‘**’ 0.01 ‘*’ 0.05.

## Notes

### Competing Interest Statement

The authors have declared no competing interest.

### Summary of Updates

This version of the manuscript has been revised to update the author list.

