## Supplementary figures and images for "Tumor-Associated Macrophages Expand Chemoresistant, Ovarian Cancer Stem-Like Cells"

### Supplemental Figure 1

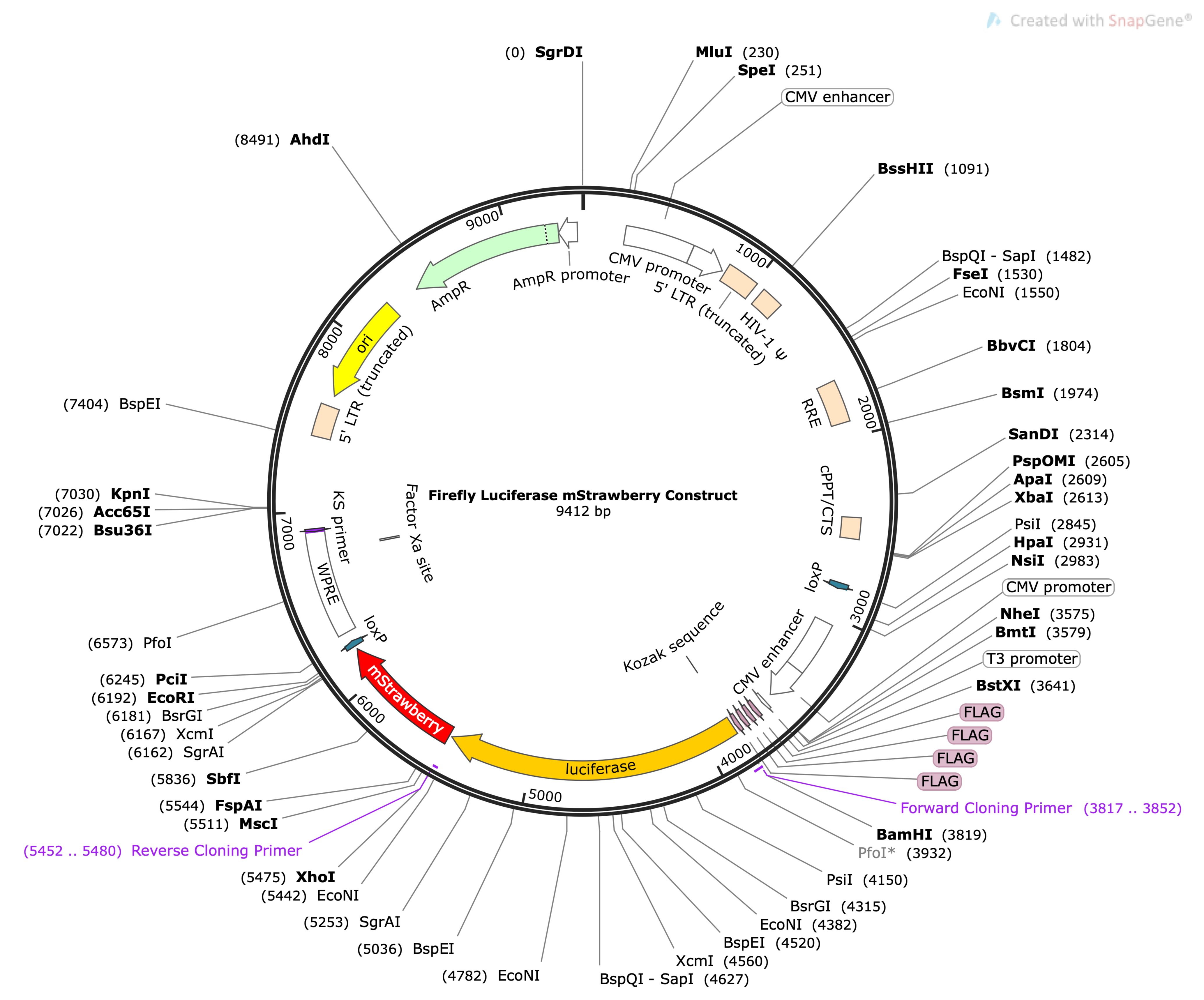

### Supplemental Figure 2

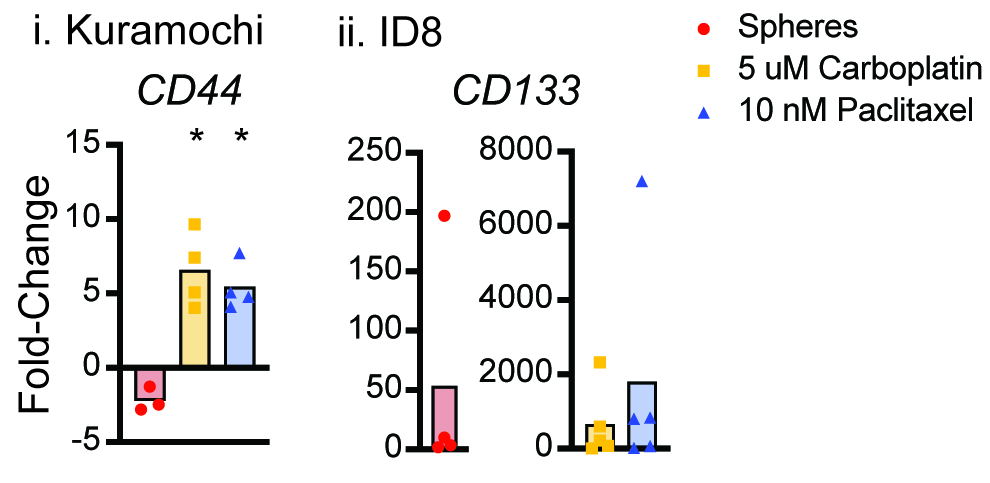

### Supplemental Figure 3

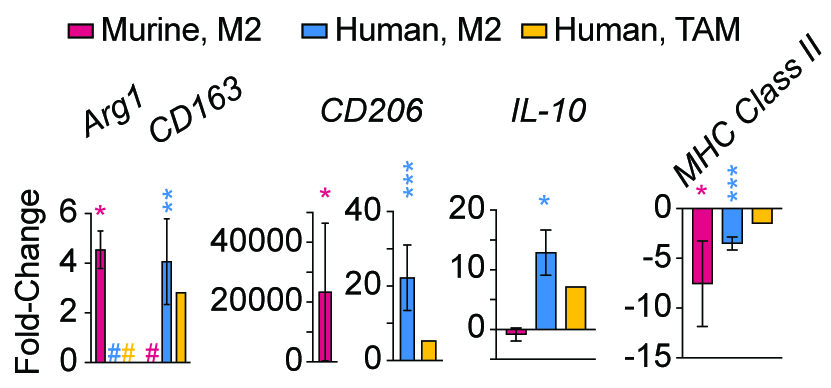
