## Supplemental Table 1 for "Tumor-Associated Macrophages Expand Chemoresistant, Ovarian Cancer Stem-Like Cells"

**Supplemental Table 1: Primers Used**

| **Gene** | **Forward Primer, Human** | **Reverse Primer, Human** | **Forward Primer, Mouse** | **Reverse Primer, Mouse** |
| --- | --- | --- | --- | --- |
| Endogenous Control | | | | |
| *Actin* | CCAACCGCGAGAAGATGA | CCAGAGGCGTACAGGGATAG | GGCTGTATTCCCCTCCATCG | CCAGTTGGTAACAATGCCATGT |
| Ovarian Cancer Stem-Like Cell Markers | | | | |
| *ALDH1A1* | CAAGATCCAGGGCCGTACAA | CAGTGCAGGCCCTATCTTCC | CAAGGTGGCCTTCACTGGAT | ATGCTGCGACACAACATTGG |
| *ALDH1A2* | AACTCTGGAACTTGGAGGCA | CTGCTCCACAGCATAGTCCA | AATCGCTTCTCACATCGGCA | CAGCGTAGTCCAAGTCAGCA |
| *ALDH1A3* | ACAAAGCCCTGAAGTTGGCT | ACAAAGCCCTGAAGTTGGCT | AGGGGCCAAGCTAGAATGTG | GTCGGTGCTATTCGCTCTCT |
| *CD24* | CACCGACGGAGGGGACAT | GTGGCATTAGTTGGATTTGGGG | CCCACGCAGATTTACTGCAAC | AGAGACGTTTCCTGGCCTGA |
| *CD44* | TGCAGTCAACAGTCGAAGAAG | GTCCTCCACAGCTCCATTGC | GGCTCTGATTCTTGCCGTCT | TGTCTTCCACCGTCCCATTG |
| *CD133* | GCCCCCAGGAAATTTGAGGA | ATCCATTCCCTGTGCGTTGA | TCTGGTGGGCTGCTTCTTTT | GGTCCCTTTGATCCGAGTCC |
| *KIT* | AACACGCACCTGCTGAAATG | GTCTACCACGGGCTTCTGTC | TTCCCGGAGCCCACAATAGA | TTGGAGGCCTTACACTCCAC |
| M2 Macrophage Markers | | | | |
| *Arg1* | n/a | n/a | GAGACGTAGACCCTGGGGAA | TTTCTTCCTTCCCAGCAGGT |
| *CD163* | ACGCGTGGAGGGAATATGTGTTCT | TCAAACCAGATTGGTCCAGAGCCT | GAGACACACGGAGCCATCAA | CGTTAGTGACAGCAGAGGCA |
| *CD206* | CCATCGAGGAAGAGGTTCGG | GGTGGGTTACTCCTTCTGCC | CTAACTGGGGTGCTGACGAG | GGCAGTTGAGGAGGTTCAGT |
| *IL-10* | CGAGATGCCTTCAGCAGAGT | CGCCTTGATGTCTGGGTCTT | CAATAACTGCACCCACTTCCC | TTTCCGATAAGGCTTGGCAAC |
| M1 Macrophage Markers | | | | |
| *IL-12* | GCACAGTGGAGGCCTGTTTA | GCCAGGCAACTCCCATTAGT | GGGACCAAACCAGCACATTG | AGATGCTACCAAGGCACAGG |
| *HLA-DR* | GAGCGCCCAAGAAGAAAATGG | GATAGAACTCGGCCTGGATGA | n/a | n/a |
| *H2-Ab1* | n/a | n/a | GCAGACACAACTACGAGGGG | GCGCACTTTGATCTTGGCTG |
| Cloning Plasmids for Firefly Luciferase Cloning into mStrawberry Plasmid | | | | |
| Forward | CGCGGATCCATGGAAGACGCCAAAAACATAAAGAAAG | | | |
| Reverse | CCGCTCGAGCAATTTGGACTTTCCGCCCTTCT | | | |
