## Supplemental Table 2 for "Tumor-Associated Macrophages Expand Chemoresistant, Ovarian Cancer Stem-Like Cells"

**Supplemental Table 2: *P* Values for Coculture Experiments in Figure 3E**

| **Comparison** | | **Rank** | ***P*  Value** | **Holm-Bonferroni *P* Cutoff** |
| --- | --- | --- | --- | --- |
| **Kuramochi** | | | | |
| No Drug,  No Macros | No Drug,  M2 | 1 | 2.45187E-05 | 0.007142857 |
| PTX,  No Macros | PTX,  M2 | 2 | 0.001326346 | 0.008333333 |
| No Drug,  No Macros | PTX,  M2 | 3 | 0.001530108 | 0.001530108 |
| Carbo,  No Macros | Carbo,  M2 | 4 | 0.001558191 | 0.0125 |
| No Drug,  No Macros | PTX,  No Macros | 5 | 0.023435 | 0.016666667 |
| No Drug,  No Macros | Carbo,  No Macros | 6 | 0.031887771 | 0.025 |
| No Drug,  No Macros | Carbo,  M2 | 7 | 0.529882095 | 0.05 |
| **ID8** | | | | |
| PTX,  No Macros | PTX,  M2 | 1 | 0.00010926 | 0.00714286 |
| Carbo,  No Macros | Carbo,  M2 | 2 | 0.00027354 | 0.00027354 |
| No Drug,  No Macros | Carbo,  No Macros | 3 | 0.0009011 | 0.01 |
| No Drug,  No Macros | No Drug,  M2 | 4 | 0.00297571 | 0.0125 |
| No Drug,  No Macros | PTX,  No Macros | 5 | 0.00667273 | 0.01666667 |
| No Drug,  No Macros | PTX,  M2 | 6 | 0.08435697 | 0.025 |
| No Drug,  No Macros | Carbo,  M2 | 7 | 0.52510097 | 0.05 |
