## Supplemental Table 3 for "Tumor-Associated Macrophages Expand Chemoresistant, Ovarian Cancer Stem-Like Cells"

**Supplemental Table 3: Coexpressed Genes in Poor Outcome TCGA and Cocultured Kuramochi Spheres**

| **50 Coexpressed Genes** | **34 with Independent Effects on Survival** | **Progression-Free Survival Hazard Ratio (Confidence Interval)** | **Logrank *P* Value** | **References if Previously Correlated to Survival** |
| --- | --- | --- | --- | --- |
| ABL2 | ABL2 | 1.51 (1.22 – 1.88) | 0.00016 |  |
| ADAM10 | ADAM10 | 1.26 (1.09 – 1.45) | 0.002 | ^42^ |
| ADAM12 | ADAM12 | 1.28 (1.09 – 1.49) | 0.002 | ^43^ |
| ADCY1 | ADCY1 | 1.29 (1.12 – 1.49) | 0.00045 |  |
| AGFG1 | ANO7 | 1.48 (1.18 – 1.86) | 0.00069 |  |
| AGPS | C5AR1 | 1.2 (1.03 – 1.39) | 0.018 | ^44^ |
| ANO7 | CD36 | 1.36 (1.18 – 1.58) | 3.7x10^-5^ | ^45^ |
| ARRDC2 | CDH23 | 1.47 (1.18 – 1.84) | 0.00068 |  |
| C5AR1 | CH25H | 1.33 (1.15 – 1.54) | 1x10^-4^ | ^46, 47^ |
| CD36 | CPA4 | 1.32 (1.14 – 1.52) | 0.00023 |  |
| CDH23 | DOCK11 | 1.65 (1.34 – 2.04) | 2.9x10^-6^ |  |
| CH25H | DUSP16 | 1.36 (1.11 – 1.68) | 0.0033 |  |
| CHUK | EPB41L3 | 1.37 (1.16 – 1.61) | 0.00015 |  |
| CPA4 | GLIPR1 | 1.53 (1.25 – 1.88) | 3.9x10^-5^ |  |
| CSNK1G1 | GRAMD4 | 1.28 (1.11 – 1.48) | 0.00084 |  |
| DERA | IL27RA | 1.25 (1.08 – 1.44) | 0.0032 |  |
| DOCK11 | ITGB1 | 1.28 (1.1 – 1.48) | 0.0014 | ^48, 49^ |
| DUSP16 | KIAA0100 | 1.21 (1.04 – 1.4) | 0.015 |  |
| EPB41L3 | LPCAT3 | 1.2 (1.01 – 1.43) | 0.034 |  |
| FEM1B | PDE1B | 1.4 (1.2 – 1.62) | 8.6x10^-6^ |  |
| GLIPR1 | PI4K2A | 1.33 (1.15 – 1.54) | 0.00011 |  |
| GNS | PREX1 | 1.42 (1.15 – 1.76) | 0.0012 |  |
| GOLGA7 | RAB8B | 1.26 (1 – 1.59) | 0.049 |  |
| GRAMD4 | RARRES1 | 1.24 (1.07 – 1.43) | 0.0034 | ^50^ |
| IL27RA | RASSF3 | 1.34 (1.05 – 1.71) | 0.019 |  |
| ITGB1 | SH3PXD2A | 1.77 (1.44 – 2.18) | 3.5x10^-8^ |  |
| KIAA0100 | SLC6A12 | 1.24 (1.06 – 1.44) | 0.0068 | ^51^ |
| LPCAT3 | SSH1 | 1.56 (1.33 – 1.83) | 2.6x10^8^ |  |
| LRP1 | STARD8 | 1.35 (1.16 – 1.58) | 0.00013 |  |
| OSBPL8 | TGFBI | 1.36 (1.15 – 1.6) | 0.00024 | ^52-54^ |
| PDE1B | TGM2 | 1.21 (1.03 – 1.42) | 0.022 |  |
| PI4K2A | TIE1 | 1.39 (1.2 – 1.61) | 7.9x10^-6^ |  |
| PPTC7 | VASP | 1.21 (1.03 – 1.43) | 0.02 |  |
| PREX1 | WNK1 | 1.34 (1.14 – 1.56) | 0.00025 |  |
| PTGER2 |  |  |  |  |
| RAB8B |  |  |  |  |
| RARRES1 |  |  |  |  |
| RASSF3 |  |  |  |  |
| S100A9 |  |  |  |  |
| SH3PXD2A |  |  |  |  |
| SLC35E3 |  |  |  |  |
| SLC6A12 |  |  |  |  |
| SSH1 |  |  |  |  |
| STARD8 |  |  |  |  |
| TGFBI |  |  |  |  |
| TGM2 |  |  |  |  |
| TIE1 |  |  |  |  |
| TMEM19 |  |  |  |  |
| VASP |  |  |  |  |
| WNK1 |  |  |  |  |
